# The Constriction Ring of Amniotic Band Syndrome Reveals Idiosyncrasies of Wound Repair in Infancy

**DOI:** 10.1101/266478

**Authors:** Surjya N. Bhattacharyya, Isaac M. Ilyashov, Alice Chu

## Abstract

We hypothesized that the constriction rings in Amniotic Band Syndrome (ABS) are the sequelae of localized mechanical injury. Typical scarring pattern was examined in skin tissue of ABS origin, containing an amniotic band constriction, for collagen and elastin distribution, and the ratio of collagen I to collagen III (CI:CIII). A skin sample from an extra finger was the control. In the ABS specimens, sub-epidermal structures were intact and present throughout, and collagen I exhibited a normal basket-weave pattern. At the site of constriction in both ABS samples, reticular dermis elastin fibers were fragmented and papillary dermis elastin fibers were absent. In the control tissue, the reticular dermis contained relatively thick, branching fibers of elastin, and papillary dermis elastin was present.

The elastin fragmentation at the constriction ring indicates localized elastin remodeling in response to injury. The absence of elastin in the papillary dermis of the constriction likely indicates a localized disruption in elastin formation. The formation and deposition of collagen and the presence of sub-epidermal structures favor a non-scarring phenotype, while the organization of elastin favors that of localized scarring.

**Summary Statement:** A rare, fetal model of healing after constrictive injury of skin, in which elastin fragmentation at the constriction injury indicates remodeling explained by the differential expression of elastin during gestation.

## INTRODUCTION

In Amniotic Band Syndrome (ABS), free floating fibrous strings within the womb called ‘amniotic bands’ wrap around lower or upper extremities or other anatomical locations. This results in the formation of fetal dermal constrictions, called ‘constriction rings.’ ABS is responsible for one in 70 stillbirths with an incidence of one in every 1,200 births (Barros et al., 2014).

Amniotic band constriction can also result in vascular compression to the affected areas of the infant, sometimes causing necrosis and auto-amputation. A case of ABS occurring with polydactyly was observed in a female infant (Robin et al., 2005). The anatomical location of these bands determines the exact deformation that appears (Barros et al., 2014). ABS occurs in association with other defects including acrosyndactyly, oligodactyly, and scoliosis (Goldfarb et al., 2009).

Two competing theories exist about the pathogenesis of ABS. The extrinsic, or exogenous, theory suggests that amniotic bands originate from amnion rupture, resulting in migration of the fetus into the chorionic cavity and subsequent compression of the fetus from close contact with the chorion (Sentilhes et al., 2003). The intrinsic, or endogenous, theory suggests amniotic bands originate during embryogenesis and formation of the amniotic cavity.^5^ Severe constriction by the bands causes fibrous scar formation that may result in vascular, lymphatic and neural damage that develops ischemia (Moran et al., 2007).

In adults, constrictive scars are the endpoint of localized tissue ischemia or inflammation (Brenner et al., 2009). Localized ischemia in wounds leads to fibrogenesis characterized by elevated collagen synthesis and remodeling (Dalton et al., 2007). In ABS, affected areas under the constriction band tend to exhibit non-normal, indented, scarring characteristics (Hajihosseini et al., 2010). Subcutaneous tissues atrophy in response to the amniotic band, resulting in a thin dermis. The type of injury seen in ABS presumably occurs from the local ischemia that band compression inflicts on the fetus within the womb (Moran et al., 2007).

Fetal skin tends to exhibit little to no scarring in response to injury, meaning that collagen is deposited in moderation, while in contrast, high collagen deposition occurs in hypertrophic scarring in adult skin. Scar-less healing in fetal skin is attributed to the absence of pro-fibrotic inflammation, which causes acute disorganization amongst reparative collagen fibers in adults. Normal fetal skin also contains a higher ratio of collagen III to collagen I, while both adolescent and adult skin contains more collagen I than collagen III (Bullard et al., 2003).

In normal skin, other components of the dermis include elastin and fibrillin-rich microfibers, which are major elements of the extra-cellular matrix (ECM). These elastic fibers form a mesh containing thick fibers within the deep or reticular dermis (Cotta-Pereira et al., 1976). The distribution of elastin has rarely been evaluated in fetal cutaneous wound healing.

The adult model of constrictive wound formation is localized ischemia leading to fibrogenic responses that affect collagen and elastin remodeling, and ABS represents a fetal analogy of the same injury (Brenner et al., 2009; Dalton et al., 2007; Baruch et al., 1997; Tsuji and Sawabe 1987). We hypothesized that the constriction rings seen in ABS represent the sequelae of localized mechanical injury with a resultant scarring pattern. We analyzed the scarring characteristics of both collagen and elastin within the dermis, as well as in the ratio of collagen I to collagen III (CI:CIII).

## MATERIALS and METHODS

### Amniotic band skin samples

ABS tissue was obtained from our IRB approved Pediatric Musculoskeletal Tissue Bank. Two samples were tested from constriction rings excised from a patient with ABS at 5 and 7 months after birth. A skin sample from an extra finger of a 20 month old was used as control tissue.

Skin tissue specimens have been surgically removed during orthopedic surgical procedures, which were performed for treatment purposes as part of standard of care. These samples, which typically would be discarded, were banked. The samples were de-identified and labeled with their unique codes.

### H&E staining & Weigart’s Resorcin fusion Staining

Basic morphology was determined via H&E staining, and elastin was stained using Weigart’s Resorcin/Fuchsin. H&E and elastin stained sections were recorded digitally using an Aperio Epathology digital imager and Imagescope software. All staining and imaging was carried out by the NYU Histopathology Core

### Picrosirius red staining

Collagen I & III were stained using picrosirius red (Junqueira et al., 1979). First, the samples were cut into 18nm sections and de-paraffinized in varying concentrations of xylene solution. The samples were then rehydrated and stained with Weigert’s Hematoxylin for 8 minutes. After rinsing the samples once more, picrosirius red was added and left to incubate for one hour, creating a near-equilibrium stain. The samples were then rinsed in two changes of an acetic acid solution. After removal of excess water with movement, they were rinsed in three changes of 100% ethanol. Lastly, the samples were cleared in xylene and mounted in resin. All picrosirius staining was carried out by the NYU Histopathology Core. Afterwards, the picrosirius stained slides were photo documented using a Leica DMLM microscope made for routine microscopy. Photos were taken in bright-field, 50% polarized light, and 100% polarized light.

Fiji image analysis software was used according to (Schindelin et al, 2012), to quantify the ratios of collagen I to collagen III in our picrosirius red stained sections. The total pixel areas of red (staining for collagen I) and green (staining for collagen III) were isolated from the picrosirius red images using Fiji’s color threshold tool. These total pixel areas were then calculated using Fiji’s analysis>measure tool. For each sample with a constriction, the collagen ratio within 8 quadrants was averaged for regions at the site of constriction and away from constriction. For the control sample, only 8 quadrants were used for calculating the collagen ratio and were categorized as away from a constriction.

Rabbit anti- LTBP1 (LifeSpan Biosciences, LS-B8256 IHC Plus) was used for immunohistochemical staining based on the manufacturer described protocol.

## RESULTS

In the ABS specimens, subcutaneous structures such as follicles, sebaceous glands, and blood vessels were fully intact and present throughout each sample (Fig. 1A,B). However, there was a 30 to 57.1% reduction in reticular dermal thickness at the site of the constriction ring, while epidermal thickness remained consistent when compared to normal skin.

**Fig. 1A:**
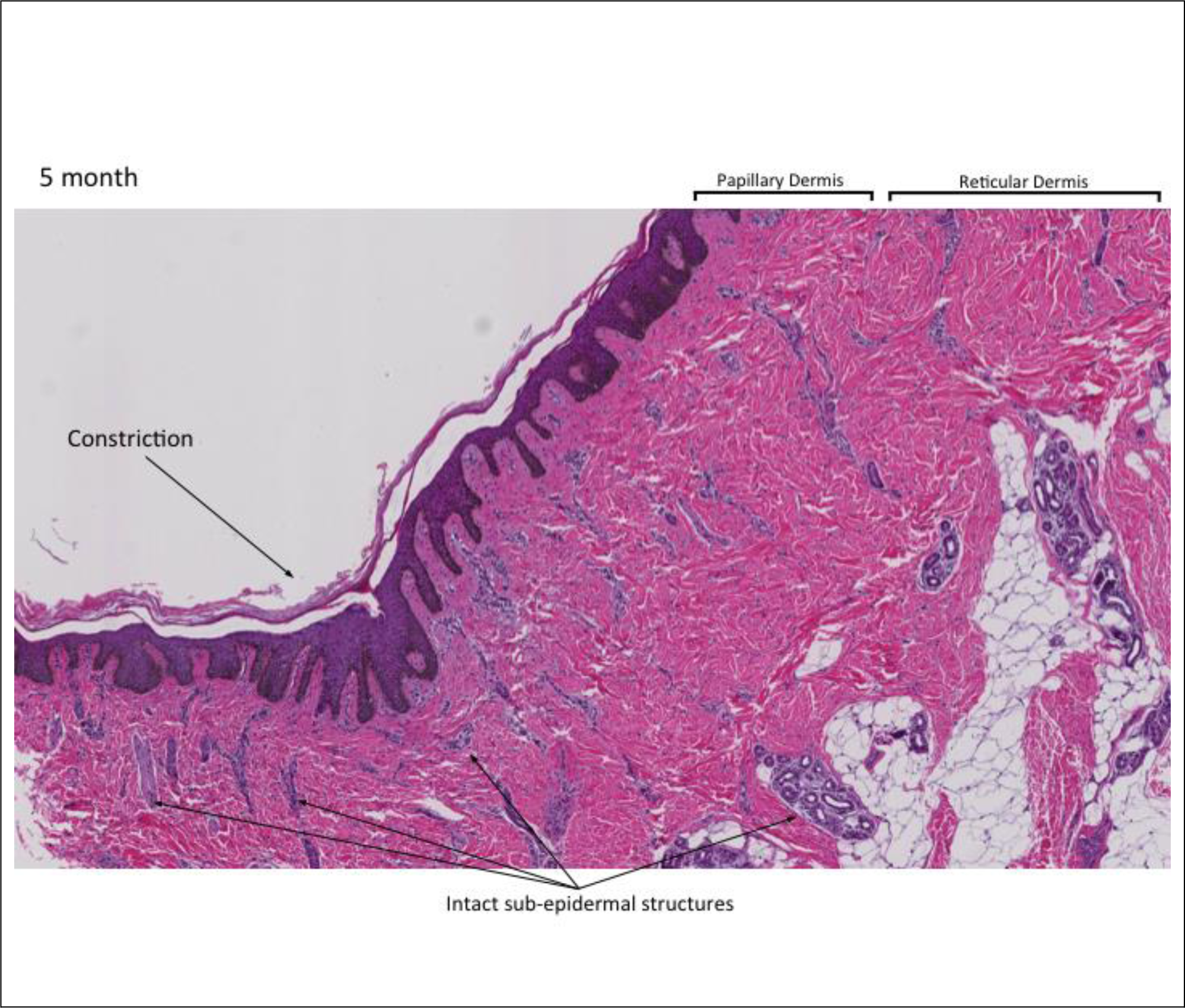
H&E staining of sample excised at 5 months. 5x magnification. Arrows indicate site of constriction and intact sub-epidermal structures.

**Fig. 1B:**
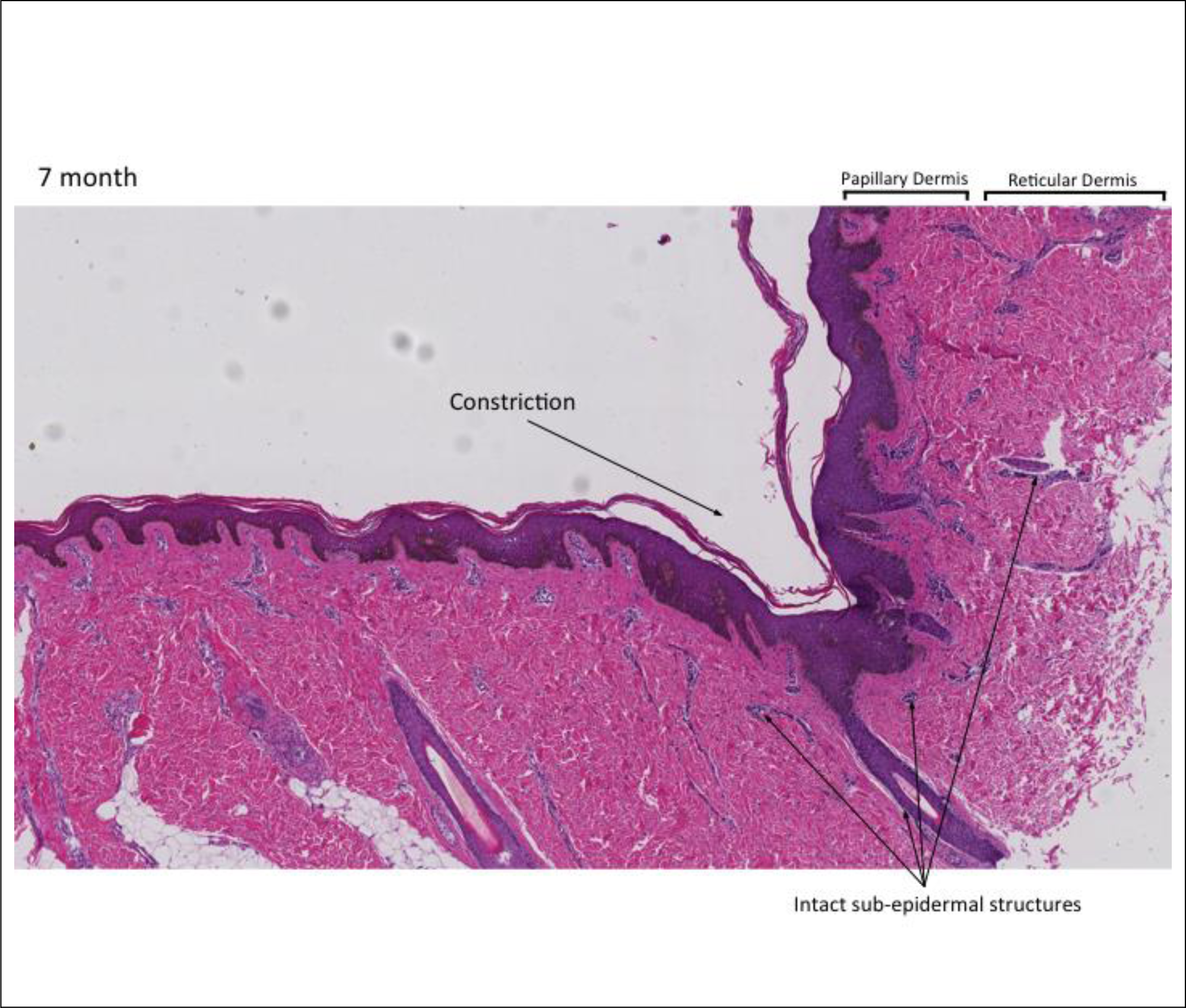
H&E staining of sample excised at 5 months. 5x magnification. Arrows indicate site of constriction and intact sub-epidermal structures.

Collagen I exhibited a normal basket-weave pattern in the picrosirius red staining of the ABS and control samples (Fig. 2A,B). We calculated the means of the ratios of collagen I: collagen III (CI: CIII) in areas below the constriction ring and away from the constriction ring as described in the methods section. Table 1 shows the mean of the ratios of collagen I: collagen III in areas below the constriction ring and away from the constriction ring. Using a two-tailed t-test for related samples, the mean CI:CIII ratios were not statistically significantly different between groups away vs. below the constriction, (Table1). The Shapiro-Wilk test for normality confirmed the data had a normal distribution.

**Fig. 2A:**
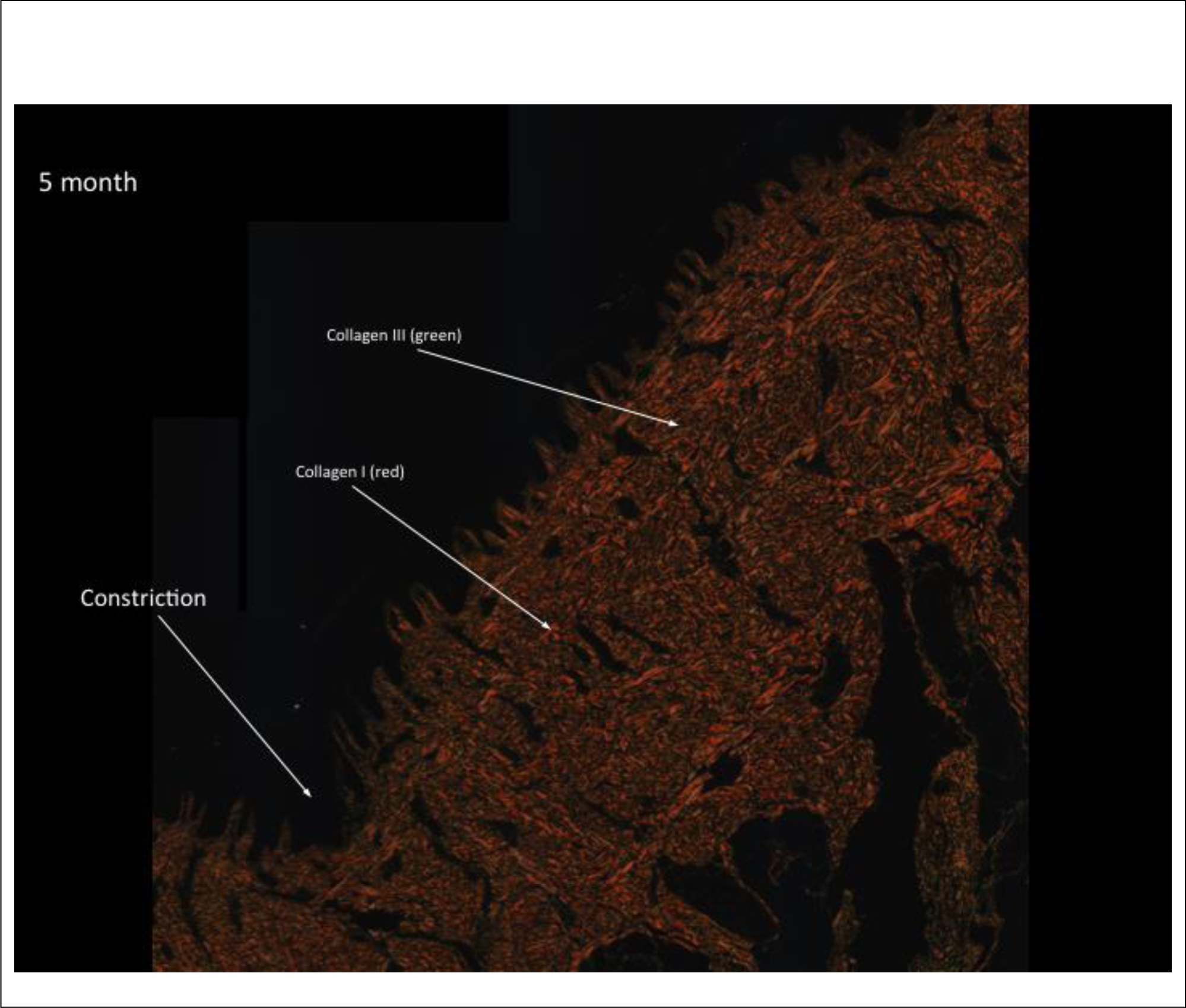
Picrosirius red staining of sample excised at 5 months. Viewed in polarized light. Collagen I is stained in red and collagen III is stained in green (faint).

**Fig. 2B:**
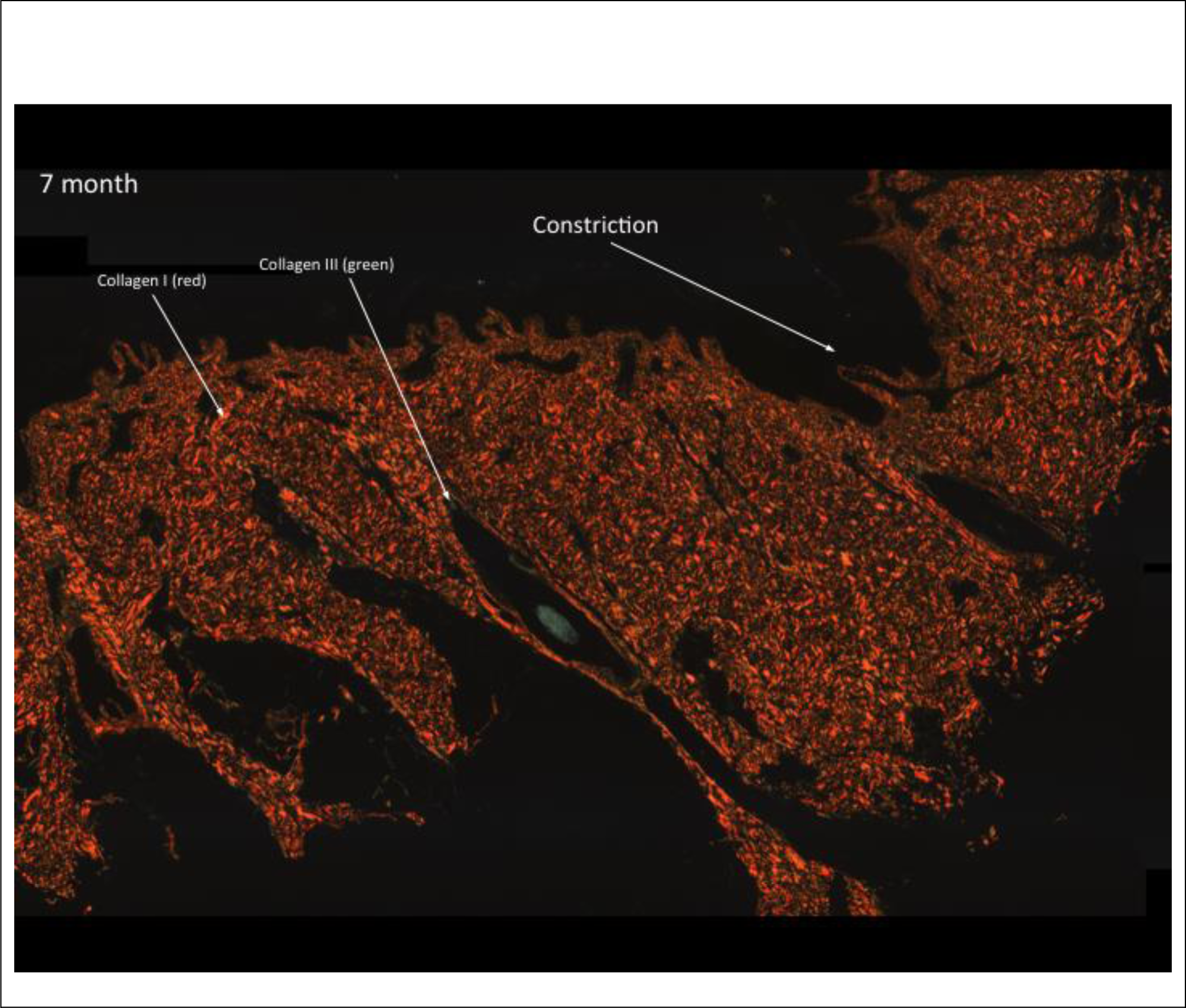
Picrosirius red staining of sample excised at 7 months. Viewed in polarized light. Collagen I is stained in red and collagen III is stained in green (faint).

**Table1:** Mean of the Ratios of Collagen I: Collagen III in samples stained with Picrosirius Red

| Location | Mean CI:CIII Ratio | P-value of grouped collagen difference |
| --- | --- | --- |
| Below Constriction | 38.2985 | 0.816 |
| Away from Constriction | 50.716 | 0.167 |

At the site of constriction in both ABS samples, reticular dermis elastin fibers were fragmented, globular, and lacked a pronounced pattern of branching. Whereas, in the control tissue, there were relatively thick, dichotomously branching fibers of elastin throughout the reticular dermis (Fig. 3). Papillary dermis elastin fibers were scant at the constriction in ABS samples compared to the control skin tissue, which displayed thin elastin fibers configured in a dichotomously branching pattern (Fig. 3).

**Fig. 3:**
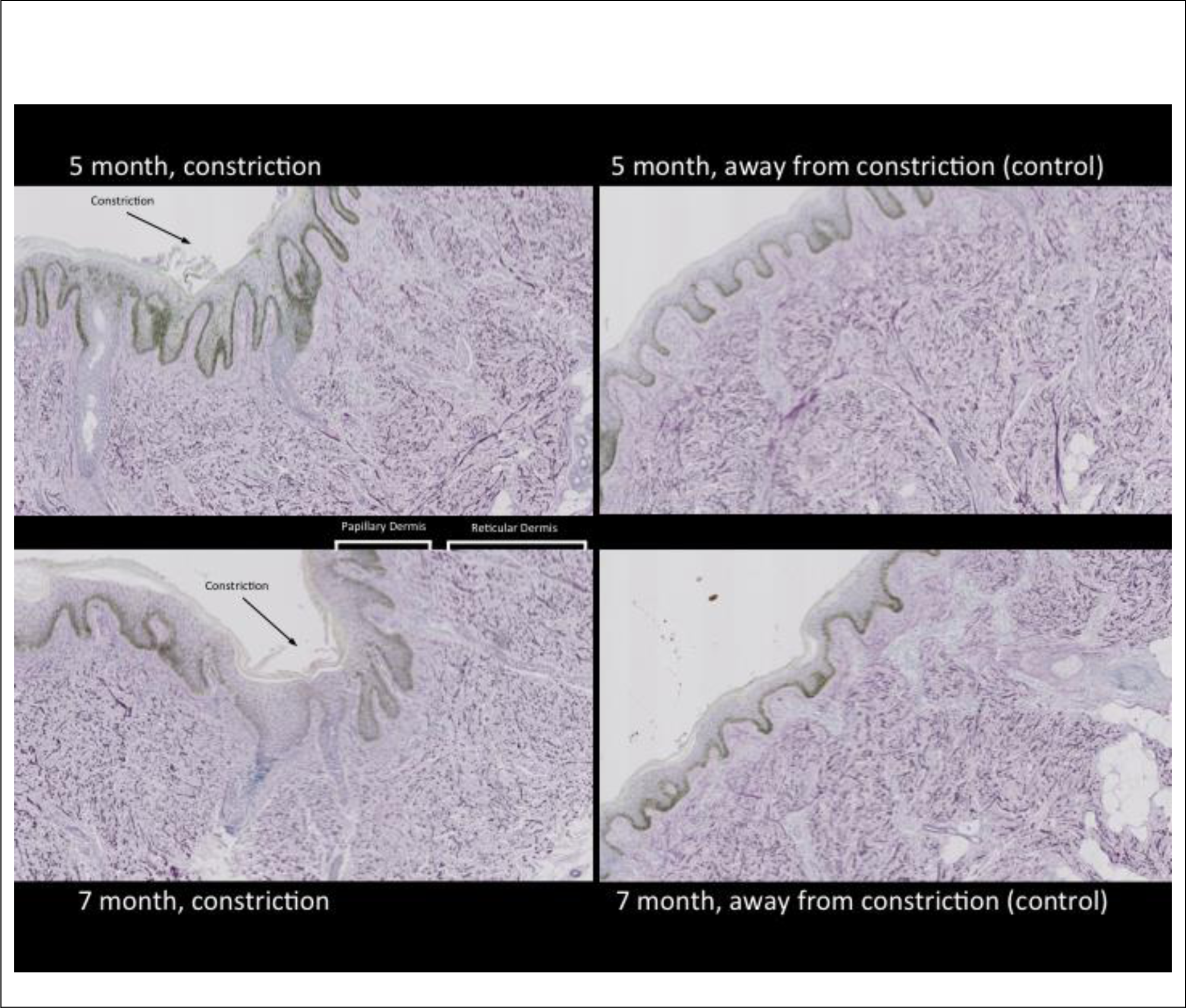
Papillary and reticular dermis regions in 7 and 5 month old, at constriction ring and away from constriction ring. Weigart’s Resorcin/Fuchsin staining showing elastin fibers stained in purple. Samples at 10x magnification.

In ABS samples (5 and 7 months year old), latent TGF binding protein (LTBP) staining was sparse throughout the dermis. Similarly, in another control skin sample from a 14 year old, LTBP staining was sparse and scattered throughout the dermis (data not shown).

## DISCUSSION

Amniotic band constriction results in dermal thinning, subcutaneous tissue atrophy, and fibrous scar formation that may cause vascular, lymphatic and neural damage (Moran et al., 2007). Subcutaneous tissues compressed by the amniotic band exhibit indented scarring characteristics (Hajihosseini et al., 2010). In adults, damaged blood vessels decrease cellular nutrition, resulting in localized ischemia, leading to wound repair, fibrogenesis, and elevated collagen synthesis and remodeling (Dalton et al., 2007; Kaplan et al., 1997).

In our ABS samples, normal sub-epidermal structures were present in the reticular dermis below the constriction ring. Since the absence of sub-epidermal structures is an important characteristic of scarring (Occleston et al., 2010; Rendl et al., 2005); the mechanical injury from the amniotic band did not create a typical scar. A non-scarring phenotype is further supported by our observations of collagen in the picrosirius stained sections. There were no significant differences in the organization of collagen fibers between the dermal regions below the constriction ring and in the surrounding areas. These collagen fibers exhibited a basket weave pattern, characteristic of normal skin (Romanelli et al., 2012; Smith et al., 1986).

Cheng et al. reported on CI:CIII ratios in fetal skin, so it was investigated whether there were differences seen in our specimens (Cheng et al., 2000). In the ABS samples, the mean CI:CIII ratios were not statistically significantly different between the groups away versus the groups below the constriction ring. It is therefore unlikely that amniotic band constriction affects the ratios of CI:CIII.

Despite the differences between wound repair in fetal and adult skin, both involve fibroblast recruitment and subsequent collagen deposition (Broughton et al., 2006). When an injury occurs, dermal fibroblasts differentiate into myofibroblasts and activate extracellular matrix (ECM) collagen production (Lijnen and Petrov, 2002; Werner et al., 2007). Both the activation of fibroblast differentiation and collagen production are caused by TGF-beta (Werner et al., 2007). In the TGF-beta pathway, TGF-beta1 is predominantly kept in a bio-inactive state within the large latent complex (LLC) bound to the ECM. The LLC contains two components: the latent TGF binding protein (LTBP) and the pro-TGF-beta1. When LTBP is cleaved from Pro-TGF-beta1, TGF-beta1 is released and wound healing commences (Annes et al., 2003; Feng and Derynck, 2005).

The sparse LTBP staining throughout the dermis, and the normal levels of collagen I at the site of the amniotic band in the ABS samples seem to indicate that TGF-beta is being expressed in large enough numbers to create the myofibroblast-produced collagen responsible for normal healing. This evidence suggests a normal regulation of collagen in ABS.

However, the excessive fragmentation and thinning seen in elastin fibers below the constriction ring does indicate the occurrence of a scarring event. Studies reported that fragmentation and thinning of elastin are characteristics of hypertrophic and atrophic scarring, respectively, in adult skin tissue (Tsuji and Sawabe, 1987; Bhangoo et al., 1976; Amadeu et al., 2004). The elastin fragmentation observed in our samples suggests hypertrophic scar qualities, while the scant distribution of elastin in the papillary dermis below the constriction ring indicates atrophic scarring. This mix of phenotypes is undocumented in the literature, but may indicate a localized disruption in elastin formation specific to fetal tissue.

Overall, our findings show a non-scarring appearance in dermal collagen, while the organization of elastin favors localized, mixed hypertrophic and atrophic scarring. This points to a disconnect between collagen and elastin deposition and their timing during fetal healing.

Amnion rupture and subsequent band formation occurs at 12 weeks of gestation, with ABS prenatally diagnosed at a median of 15 weeks (Morovic et al., 2004; Barros et al., 2014). Normal collagen deposition occurs during 0-24 weeks of gestation in the womb and is characterized by scarless fetal skin healing (Bullard et al., 2003; Lorenz et al., 1995). Due to the timing of these events, collagen fibers at the constriction rings would have a 12-week period between the 12th to 24th weeks of gestation to undergo repair with characteristics of scarless healing (Fig. 4). This may explain why the collagen fibers were observed to have undergone normal deposition (Fig 2), and that sub-epidermal structures are present and intact in the ABS samples (Fig. 1A,B).

**Fig. 4:**
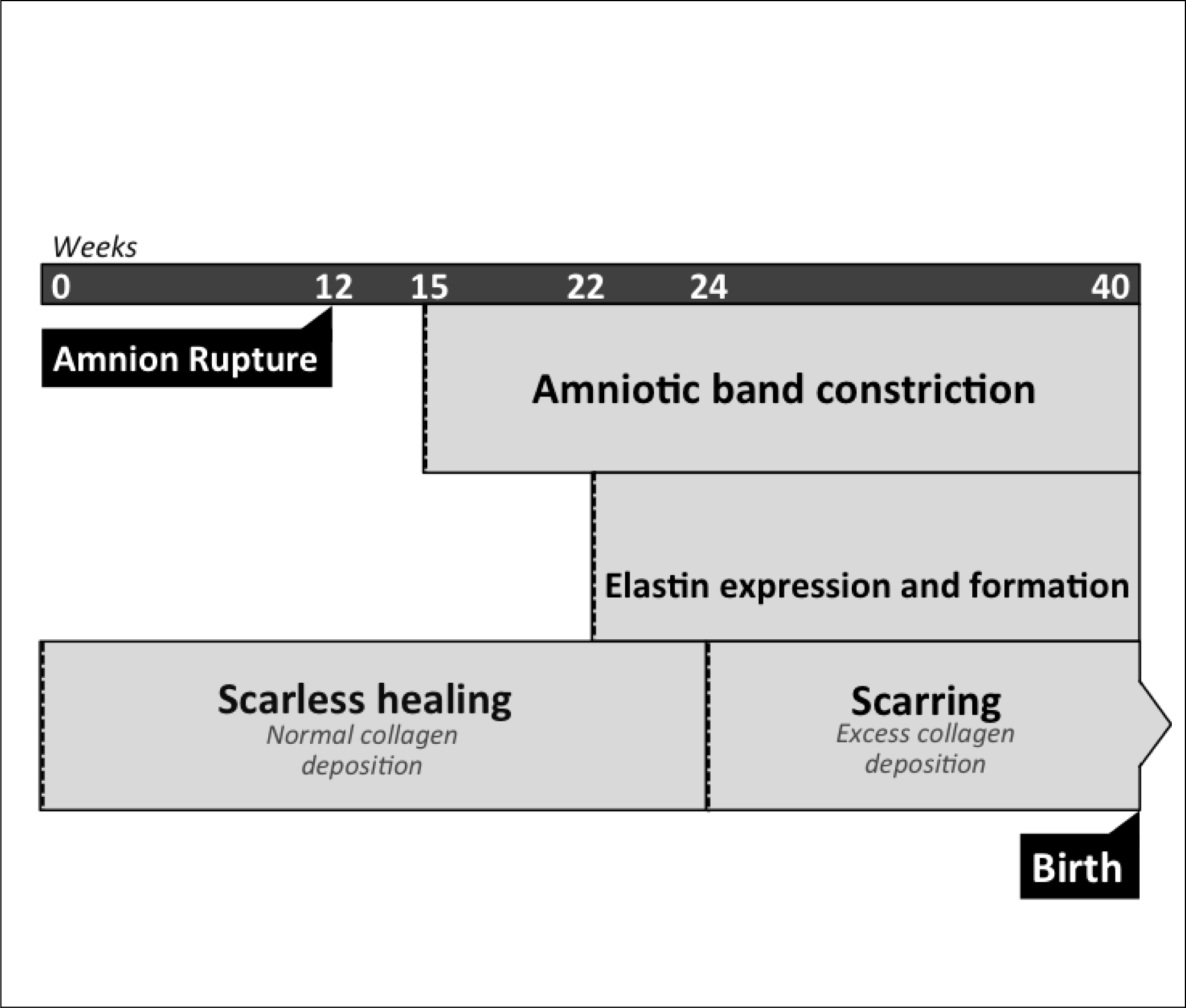
Temporal developments during gestation. Amnion rupture and subsequent band formation occurs at 12 weeks of gestation. The amniotic band constriction develops at a median of 15 weeks. Scarless fetal skin healing occurs during 0-24 weeks of gestation in the womb when there is normal collagen deposition. The expression and formation of elastin occurs from 22-40 weeks of gestation.

The expression and formation of elastin occurs from 22-40 weeks of gestation (Fig. 4) (Deutsch and Esterly, 1975; Sephel et al., 1987; Holbrook and Smith, 1981). Since the ABS constriction exists before the expression of elastin, elastin fibers have 18 weeks to develop and undergo repair while under amniotic-band-induced mechanical stress. This correlates with our observation of localized disruption in elastin formation in ABS (Fig. 3). Our work suggests that ABS scarring is because of an inability to remodel dermal elastin later in gestation.

There are several proteins of interest that are correlated with elastin development. Fibrillin-1 is vital in elastin deposition during development. In conjunction with fibroconectin, fibrillin-1 provides microfibril scaffolding for the formation of elastin fibers. Localized fibroblasts produce fibrillin-1, which binds to heparan sulfate on cell surfaces in the form of “bead-on-string” microfibrils. These microfibrils later aggregate around fibrocenectin in the ECM and form an extracellular microfibril network (Sabatier et al., 2014). Fibroblasts also produce elastin precursor tropoelastin, which binds to heparan sulfate on cell surfaces in the form of tropoelastin globules. Fibulin-5 complexes with these tropoelastin globules and transfers them to the extracellular fibrillin microfibril network, where they combine into strands of elastin (Sideek et al., 2014). Mutations in the fibrillin-1 gene FBN1 have been linked to several connective tissue disorders, due to the ECM and bone related complications that arise from altered fibrillin expression, such as in Marfan’s syndrome (Lee and Godfrey, 1991; Loeys et al., 2001; Ramachandra et al., 2015).

The conflicting evidence of wound healing seen in ABS likely represents a skewed scarring phenotype, with collagen and elastin healing differently due to the temporal uniqueness of fetal healing. It is likely that the scarless nature of fetal healing does not apply to elastin remodeling. Further research is required to elucidate the intricacies of fetal wounding and the role of elastin during the development of ABS.

In summary, ABS represents a rare, fetal model of healing after constrictive injury. Our initial hypothesis was that ABS would be analogous to known adult models in terms of collagen deposition and elastin morphology. Surprisingly, dermal collagen was intact, consistent with the scarless nature of fetal skin healing. However, dermal elastin was disrupted, and we believe that to be explained by the differential expression of elastin during gestation. These findings are a unique and previously unreported observation of fetal response to mechanical tissue injury.

## Acknowledgements

Histopathology Core, NYU School of Medicine.

David Feldman MD, Paley Institute.

Andrew Myles Parrott, Department of Pathology and Cell Biology, Columbia University Medical College.

## Competing interests

The authors declare no competing interests.

## Funding

This work was supported by funds from KiDS of New York University Langone.

